## Supplementary material for "Defining expansions and perturbations to the RNA polymerase III transcriptome and epitranscriptome by modified direct RNA nanopore sequencing": Table S1

| **transcript** | **chr** | **start** | **end** | **strand** | **sequence (5’ -> 3’)** |
| --- | --- | --- | --- | --- | --- |
| tRNA-Glu-CTC-2-1 | 1 | 248,874,246 | 248,874,338 | + | CTTccctggtggtctagtggttaggattcggcgctctcaccgccgcggcccgggttcgattcccggtcaggaaaGTAAGCCGTTTT |
| tRNA-Thr-CGT-4-1 | 17 | 31,550,062 | 31,550,226 | + | CTGGGCTGTCAGGCGCGGTGGCCAAGTGGTAAGGCGTCGGTCTCGTAAACCGAAGATCGCGGGTTCGAACCCCGTCCGTGCCTGAGACCCGAGGTAGGGCTTTGGCTGTGGGGAAGTCGGGTTTTCTCCACGTACGCCGTCCCTTCTACGTGGCATTTT |
| tRNA-Pro-TGG-1-1 | 14 | 20,633,008 | 20,633,112 | + | CTCGTTGGTCTAGTGGTATGATTCTCGCTTTGGGTGCGAGAGGTCCGGTTCAATCCCGGACGAGCCCTTACTTTCCTTTCCGTTTCATCTTTCTCTCTTT |
| tRNA-Arg-CCT-3-1 | 16 | 3,152,898 | 3,152,994 | + | gccccggtggcctaatggataaggcattggcctcctaagccagggattgtgggttcgagtcccacccgggtaAAGAAGGCCGAATTTT |
| novel tRNA 1 | 11 | 68,460,132 | 68,460,222 | - | CTAGGGAATCAGCTTAAGTGGAGGAGCGTTCGTTTAGTATGTGAGAGGTAACGGGATCGATGCCTGCATTCTCCACAGGGGAATTTT |
| novel tRNA 2 | 1 | 22,255,657 | 22,255,795 | - | ATTTACAGTCCCGGCACTGTGGCTCACGCCTATAATCCCAGCAATTTGGGAGGCCAAGGAGGCAGGATCACTTGGGGCCAAGAGTTCAAGACCAGTTTGGCCAACATAGTAAGACCCTGTCTCTATTTAATACATTTT |
| novel tx 1 / AluJb SINE Alu | 17 | 17,960,225 | 17960341 | + | ACTAAGCTCTCtggtttagtggttaagagcaccagctctgctgcccacagacctgggttcaatccctgctctgccactgataatccacagaccttgaggaaggtacttgactttt |
| novel tx 2 / nc111 | 1 | 160,925,972 | 160,926,060 | - | gcctgatgctgtggcttagtggataagactctgtcttttcacagtggtggccagggttcaattcccgacttagggaatgagtactttt |
| novel tx 3 | 7 | 18,038,540 | 18,038,668 | + | TAATTGCAGAGGGGcagtttggtgttgtggatatgcacacagactctggtgccagactgcctggtttgaatcctggctcattaacctaagatctgggtgacttggggcaaattacttaacacatttt |
| novel tx 4 | 10 | 49,387,375 | 49,387,496 | + | TGggtcaggcgtggtggctcaggcatgcagtcccaggactttcggatgccaaggtgggcggatcacttgaggttgggagtttgagaccagcctggccaacatgatgaaacccctctctttt |
| novel tx 5 | 7 | 28,405,584 | 28,405,720 | - | TCAAggccaggcgcagtggctcacacctgtaaccacagctctttgggaggctgagacgggaggataacttgagcctgggagttcgagaccagcttgggcaacatagacagcctgagcgtctctacaaaaagttttt |
| novel tx 6 | 8 | 106,864,056 | 106,864,155 | - | ggccgggcgtggtggctcaagcctataatcccagcactttggtaagcagaggtgggtggatgacaaggtcaggagtttgagaccagcagtggccaCTTTT |
| novel tx 7 | 16 | 22,298,375 | 22,298,625 | - | TAACGGcagtatggttaagtgggtaagagcttggacccgagaacaaaactgtgcaggttcaaatcccaccacagctgcttggcatctgtgacccacctgagttgtctgggtttggtattcctccctattttcatccctaaagtagggaaactaagtaccgacctcCAGACCCCTGCGGGGAAGGAGTAAGGACAGGACGCTAATAAACGTAACCTCTGGGAAGGTTTGTTATTACTTGACAACATGTTTT |
| novel tx 8 | 12 | 93,243,600 | 93,243,720 | - | ctgagagccTgtggtctagaggagaaacacggacttgggagtagacagacatgggttctggtaccagttcttccacttccggtgtgcctttgagagagctgcttctttgagctttggtttt |

**Supplementary Table 1:** novel transcripts identified by DRAP3R
