## Supplementary material for "Defining expansions and perturbations to the RNA polymerase III transcriptome and epitranscriptome by modified direct RNA nanopore sequencing": Table S2

| **ID** | **Sequence 5’ –> 3’:** |
| --- | --- |
| qPCR primers | |
| 18srRNA_PF | AGGAATTGACGGAAGGGCACCA |
| 18srRNA_PR | TTATCGGAATTAACCAGACAAATCG |
| RMRP_F | AAGAAGCGTATCCCGCTGAG |
| RMRP_R | GCACTGCCTGCGTAACTAGA |
| RN7SK_qF | CCAGGGTTGATTCGGCTGAT |
| RN7SK_qR | GATGGTCGTCCTCTTCGACC |
| Novel tx_1_F | CTCTCTGGTTTAGTGGTTAAGAGCA |
| Novel tx_1_R | AGGTCTGTGGATTATCAGTGGC |
| Novel tx_2_F | AAAGTACTCATTCCCTAAGTCGG |
| Novel tx_2_R | CTGATGCTGTGGCTTAGTGGATA |
| Novel tx_3-new_F | TGCCAGACTGCCTGGGTTT |
| Novel tx_3-new_R | TAAGTAATTTGCCCCAAGTCACC |
| Novel_tx_4_F | ATGCAGTCCCAGGACTTTCG |
| Novel_tx_4_R | CAGGCTGGTCTCAAACTCCC |
| Novel_tx_5_F | CAGGCTGTCTATGTTGCCCA |
| Novel_tx_5_R | GGCTCACACCTGTAACCACA |
| Novel_tx_6-new-F | GCCACTGCTGGTCTCAAACT |
| Novel_tx_6-new-R | CGTGGTGGCTCAAGCCTATAA |
| Novel_tx_7_F | AGCGTCCTGTCCTTACTCCT |
| Novel_tx_7_R | GACCCACCTGAGTTGTCTGG |
| Novel tx_8-new_F | AAGCAGCTCTCTCAAAGGCA |
| Novel tx_8-new_R | ACGGACTTGGGAGTAGACAGA |
| POLR3A_F | ATGGTGAAGGAGCAGTTCCG |
| POLR3A_R | CATCCTATGGTCGAGCACCC |
| c-MYC-F | TACAACACCCGAGCAAGGAC |
| c-MYC-R | CTAACGTTGAGGGGCATCGT |
| RNA5-8SN5-F | CTTAGCGGTGGATCACTCGG |
| RNA5-8SN5-R | AGTGCGTTCGAAGTGTCGAT |
| VTRNA1-1-F | TGGCTTTAGCTCAGCGGTTA |
| VTRNA1-1-R | GGGTCTCGAACAACCCAGAC |
| RNU2-1-F | ATCGCTTCTCGGCCTTTTGG |
| RNU2-1-R | CCTATTCCATCTCCCTGCTCC |
| IVT template generation primers | |
| EBER2-ivt_F | TATTAGTACTTAATACGACTCACTATA AGGACAGCCGTTGCCCTA |
| EBER2-ivt_R | AAAATAGCGGACAAGCCGAA |
| RN7SK-ivt-F | TATTAGTACTTAATACGACTCACTATA GGATGTGAGGGCGATCTGG |
| RN7SK-ivt-R | TTGGATGTGTCTGGAGTCTTGG |
| **DRAP3R Reverse Transcription Adaptor** | |
| Oligo A | /5PHOS/GGCTTCTTCTTGCTCTTAGGTAGTAGGTTC |
| Oligo B-TN | GAGGCGAGCGGTCAATTTTCCTAAGAGCAAGAAGAAGCCAAAANN |

**Supplementary Table 2:** List of primers and oligonucleotides used in this study
